# Pannexin 1 crosstalk with the Hippo pathway in malignant melanoma

**DOI:** 10.1101/2024.09.03.611059

**Authors:** Samar Sayedyahossein, Kenneth Huang, Christopher Zhang, Mehdi Karimi, Mehrnoosh Bahmani, Brooke L. O’Donnell, Brent Wakefield, Zhigang Li, Danielle Johnston, Stephanie E. Leighton, Matthew S. Huver, Lina Dagnino, David B. Sacks, Silvia Penuela

**Author notes:** To whom correspondence should be addressed: Silvia Penuela, Department of Anatomy and Cell Biology, University of Western Ontario, London, Ontario, Canada.

## Abstract

In this study, we explored the intricate relationship between Pannexin 1 (PANX1) and the Hippo signaling pathway effector, Yes-associated protein (YAP). Analysis of The Cancer Genome Atlas (TCGA) data revealed a significant positive correlation between PANX1 mRNA and core Hippo components, YAP, TAZ, and Hippo scaffold, IQGAP1, in invasive cutaneous melanoma and breast carcinoma. Furthermore, we demonstrated that PANX1 expression is upregulated in invasive melanoma cell lines and is associated with increased YAP protein levels. Notably, our investigations uncovered a previously unrecognized interaction between endogenous PANX1 and the Hippo scaffold protein IQGAP1 in melanoma cells. Moreover, our findings revealed that IQGAP1 exhibits differential expression in melanoma cells and plays a regulatory role in cellular morphology. Functional studies involving PANX1 knockdown provided compelling evidence that PANX1 modulates YAP protein levels and its co-transcriptional activity in both melanoma and breast carcinoma cells. Importantly, our study showcases the potential therapeutic relevance of targeting PANX1, as pharmacological inhibition of PANX1 using selective FDA-approved inhibitors or PANX1 knockdown reduced YAP abundance in melanoma cells. Furthermore, our Clariom™ S analysis unveiled key genes implicated in cell proliferation, such as neuroglin1 (NRG1), β-galactoside binding protein, galectin-3 (LGALS3), that are affected in PANX1- deficient cells. In summary, our investigation delves into the intricate interplay between PANX1 and YAP in the context of invasive melanoma, offering valuable insights into potential therapeutic strategies for effective treatment.

## Introduction

Cutaneous melanoma accounts for 1.7% of newly diagnosed cancers and 0.7% of deaths among all cancers^1^. In the USA, melanoma accounted for 72% of all skin cancer-related deaths in 2017,^2^. While being the least prevalent form of skin cancer, melanoma is the most fatal, resulting in approximately 60,000 global deaths annually^3^. The transformation of melanocytes to melanoma is a result of complex intrinsic and extrinsic factors, as well as immune-related factors^4^. Mutations of signaling effectors BRAF (50%), neuroblastoma RAS viral oncogene homolog (NRAS) (26.4%) and neurofibromatosis 1 (NF1) (14.3%) are prevalent in patients with malignant melanoma^3,5^. Mutated BRAF exert its oncogenic effects through activation of the mitogen-activated protein kinase (MAPK) pathway. MAPK is the most commonly altered pathway in malignant melanoma, which resulted in mitogen-activated protein kinase kinase (MEK)-targeted therapies^6,7^. While patients respond to the combination of B-Raf/MEK inhibition therapies in the initial phases of treatment, resistance to therapy develops in the majority of patients^8^. Despite the advent of immunotherapies, 40% of patients do not respond to current treatment modalities^5,9^, highlighting the need for new treatment interventions. A role for the Hippo pathway in deriving melanoma invasion has recently emerged^10^. Yes-associated protein (YAP) is a major transcriptional co-activator of the Hippo pathway^11^. YAP transcriptional activity increases in invasive melanoma cells both *in vitro* and *in vivo*^12,13^. Importantly, increased YAP activity is required for melanoma to switch from a proliferative to invasive state^12^.

Pannexin 1, a member of a glycoprotein family (PANX 1, 2 and 3), form oligomers to establish large pore channels between the intracellular and extracellular space for cell communication^14^. Among pannexins, PANX1 has been the primary focus of research because of its widespread expression^15^. PANX1 mediates the release of small signaling molecules, such as adenosine tri- phosphate (ATP)^16^. Additionally, intracellular PANX1 has been reported to function as a calcium leak channel in the endoplasmic reticulum^17,18^. PANX1 plays an important role in normal physiological processes, including skin development and wound healing, as well as in pathological disorders, including Alzheimer’s disease, diabetes, inflammation, and cancer^14,19^. There is growing interest in investigating the role of PANX1 in the regulation of cellular processes, such as proliferation, migration, differentiation, and invasion^20,21^, which are important during melanoma tumorigenesis. Currently, our understanding of the mechanisms through which PANX1 regulates cellular processes is limited. In this paper, we investigated the possible crosstalk between PANX1 and components of the Hippo pathway in malignant melanoma. Our findings indicate that PANX1 mRNA correlates with mRNA of key components in the Hippo pathway. Furthermore, endogenous PANX1 associates with IQGAP1, a Hippo signaling scaffold^22^, in melanoma cells. In addition, knocking down or selective inhibition of PANX1 with Food and Drug Administration (FDA)-approved repurposed drugs effectively decreases the abundance of YAP and reduces YAP transcriptional activity in cells. Overall, our data indicate that PANX1 modulates YAP and can be targeted in novel therapeutic approaches against invasive melanoma.

## Methods

### *In silico* analysis of correlation between PANX1 and YAP, TAZ and IQGAP1 protein expression in melanoma and breast carcinoma

mRNA expression z-scores (RNA Seq V2 RSEM) for PANX1, YAP, TAZ and IQGAP1 were generated using data in cBioPortal.org from the Skin Cutaneous Melanoma and Breast Invasive Carcinoma Cohorts generated by TCGA Research Network (http://cancergenome.nih.gov).

### Cell lines and culture conditions

Human melanoma cell lines were obtained from American Type Culture Collection (ATCC) (Manassas, VA, USA), while the melanoma cell line 131/4-5B1 was a gift from Robert S. Kerbel^23^. Rat1 fibroblasts expressing constitutively active H-RasV12 (27) were generously provided by Marc Symons (Picower Institute, NY)^24^. Cells were cultured in Dulbecco’s Modified Eagle Medium 1X (DMEM 1X) containing 4.5g/L D-glucose, L-glutamine, 110mg/L sodium pyruvate, 10% fetal bovine serum (FBS, Invitrogen^TM^, USA), 100 units/mL penicillin, and 0.1 mg/mL streptomycin at 37°C with 5% CO_2_. Trypsin (0.25%, 1mM EDTA 1X; Life Technologies) was used to dissociate cells from culture dishes. WM239A and WM852 cells were grown in MCDB153 medium supplemented with 20% L15, 2% fetal bovine serum and 5 μg/ml insulin.

### Protein extraction and immunoblotting

Protein lysates were extracted with 1% Triton X-100, 150mM NaCl, 10mM Tris, 1mM EDTA, 1mM EGTA, and 0.5% NP-40 or RIPA buffer (50mM Tris-HCl, pH 8.0, 150mM NaCl, 1% NP- 40 (Igepal), and 0.5% sodium deoxycholate). Each buffer contained 1 mM sodium fluoride, 1 mM sodium orthovanadate, and half of a tablet of complete-mini EDTA-free protease inhibitor (Roche, Mannheim, Germany). Protein was quantified by the bicinchoninic acid assay (Thermo Fisher Scientific). Protein lysates (40μg) were separated by 10% SDS-PAGE and transferred onto a nitrocellulose membrane using an iBlotTM System (Invitrogen, USA). Membranes were blocked with 3% bovine serum albumin (BSA) with 0.05% Tween-20 in 1X phosphate buffer saline (PBS), and incubated with anti-human PANX1 (1:1000; PANX1 CT-412; 0.35μg/μL)^25^, anti-phospho YAP S127 (Cell Signaling # 13008), or anti-YAP (Cell Signaling # 14074) antibodies. Anti-IQGAP1 polyclonal antibodies have been characterized previously^26^. Loading controls were probed with anti-GAPDH antibody (1:1000; Millipore Cat# MAB374). For detection, IRDye® −800CW and −680RD (Life Technologies^TM^, USA) were used as secondary antibodies at 1:10,000 dilution and imaged using a LI-COR Odyssey infrared imaging system (LI-COR Biosciences, USA). Western blot quantification was conducted using Image Studio™ Lite (LI-COR Biosciences).

### PANX1 inhibitors

Carbenoxolone disodium salt (Sigma Aldrich), water-soluble Probenecid (77mg/mL; Invitrogen) and DMSO-soluble Spironolactone (SelleckChem) were dissolved in Hanks’s Balanced Salt Solution (HBSS 1X, Life Technologies; calcium chloride, magnesium chloride and magnesium sulfate) to develop stock concentrations of each compound.

### Generation of CRISPR/Cas9 knockout cells

PANX1 knockout cells were generated using CRISPR/Cas9 D10A following the Ran *et al*. protocol^27^. Briefly, cells were transfected with 1 ug each of pSpCas9n(BB)-2A-Puro (PX462) V2.0 and pSpCas9n(BB)-2A-GFP (PX461) (Addgene.org) containing guide RNA sequences for human PANX1 in a 6-well plate. PANX1 gRNAs were designed with http://tools.genome-engineering.org (sequences GTTCTCGGATTTCTTGCTGA and CTCCGTGGCCAGTTGAGCGA). 24 h post transfection, cells were selected with 1 ug/ml puromycin for 72 h. Following selection, cells were screened for PANX1 levels by Western blot. Addgene plasmids were a gift from Feng Zhang (Addgene plasmid #48140 and #62987).

### shRNA knockdown of PANX1

A375-P cells were transfected with two shRNA constructs (PANX1 shRNA-B and PANX1 shRNA-D) from Origene PANX1 human 29-mer shRNA kit in pRS vector (#TR302694) (sequence: 5’- CGCAATGCTACTCCTGACAAACCTTGGCATGTCAAGAGCATGCCAAGGTTTGTCAGGAGTAGCATTGTT-3’) plus a GFP shRNA cassette (5’GCCCGCAAGCTGACCCTGAAGTTCATTCAAGAGATGAACTTCAGGGTCAGCTTGCT TTTT-3’) from Addgene (#30323) as a control. Cells transfected with control shRNA were evaluated for PANX1 expression to compare to non-transfected cells. PANX1 shRNA-expressing cells from two constructs (B and D) were selected with puromycin and examined for PANX1 knockdown. Stable knockdown samples showed 80-90% reduction in PANX1 expression. Cells were maintained under puromycin selection pressure and periodically examined for effective PANX1 knockdown by Western blot. Experiments were conducted after verifying at least 70– 85% knockdown of PANX1 protein levels by Western blotting.

### Constructs and transfection

The construction of GFP-tagged IQGAP1 has been previously described^28^. In brief, full-length IQGAP1 was tagged at the N terminus with tandem enhanced green fluorescent protein (EGFP) and inserted into pEGFP-C1 (Clontech) at a BglII and SmalI site, producing the plasmid pEGFP- IQGAP1. Stable cell lines were selected with 400 ug/mL G418. Myc-tagged wild type human IQGAP1 in a pcDNA3 vector was used^29^ ^26^. The construction of IQGAP1ΔGRD has been described previously^30^. For transfection, cells were cultured and transfected essentially as previously described^31,32^. Briefly, cells were grown in Dulbecco’s modified Eagle medium (DMEM) supplemented with 10% (v/v) fetal bovine serum. Cells were transiently transfected with 10 µg pcDNA3 (empty vector), myc-IQGAP1 or mycIQGAP1ΔGRD using FuGENE 6.

### [^3^H]thymidine uptake

Rat1 or Rat1RasV12 (which stably overexpress constitutively active H-Ras) cells were plated on 24-well culture dishes and transiently transfected with either vector control (V), wildtype (WT) IQGAP1 or IQGAP1ΔGRD (ΔGRD). After 24 h in DMEM/0.5% serum, 1 μCi/ml [^3^H]thymidine was added for 18 h. Cells were washed with ice-cold 0.5% trichloroacetic acid, lysed in 0.25 M NaOH and [^3^H]thymidine in 600 μl of lysate was quantified in a liquid scintillation counter.

### Immunoprecipitation

131/4-5B1, A375-P or A375-MA2 cells were plated in 10-cm dishes to reach 80% confluence. The following day, the cells were washed with ice-cold PBS and lysed with 500 µl of Buffer A (50mM Tris-HCl, pH 7.4, 150mM NaCl, and 1% Triton X-100) with either 1mM CaCl_2_ or 1mM EGTA supplemented with complete protease and phosphatase inhibitors (Roche, Mannheim, Germany). Lysates were subjected to two rounds of sonication for 10 s each, and insoluble material was precipitated by centrifugation at 13,000 x g for 10 min at 4 °C. Supernatants were precleared with protein A-Sepharose beads for 1 h. Equal amounts of protein lysate were incubated with protein A-Sepharose beads and anti-PANX1 polyclonal or anti-IQGAP1 polyclonal antibodies overnight at 4 °C. Rabbit IgG was used as control. Samples were washed five times with Buffer A, resolved by SDS-PAGE and Western blotting. Quick Western detection kit (LI-COR Biosciences, #926-69100) was used to probe for Pannexin 1.

### Quantitative RT-qPCR

To measure mRNA, cells were cultured for 24 h. Then total RNA was isolated from the cells using an RNA isolation kit (Qiagen). 250 ng of RNA was reverse transcribed to cDNA using a high-capacity cDNA reverse transcriptase kit (Applied Biosystems, #4374966) according to the manufacturer’s instructions. RT-qPCR was performed using SYBR Green PCR Master Mix (BioRad#1725274) and 200 nM forward and reverse primers. The primers used were human CTGF: 5’-AAAAGTGCATCCGTACTCCCA-3’ (forward), and 5’- CCGTCGGTACATACTCCACAG-3’ (reverse). RT-qPCR enzyme activation was initiated for 10 min at 95 °C and then amplified for 40 cycles of a two-step PCR (15 s at 95 °C and 1 min at 60 °C). All samples were assayed at least in duplicate, and YWHAZ was used as reference control. The results were analyzed using the ΔΔCT method.

### Immunofluorescence microscopy

Cells were grown on glass-coverslips and were fixed 72 h post-transfection using ice-cold 8:2 methanol:acetone for 15 min at 4°C, and blocked with 2% BSA-PBS. Coverslips were incubated with anti-human PANX1 antibody (1:500; PANX1 CT-412; 0.35 μg/μL) and anti-human IQGAP1 (1:100, ThermoFisher# 33-8900). Hoechst 33342 (1:1000), and Alexa Fluor 555 goat anti-rabbit IgG (2 mg/mL, 1:500) secondary antibodies, and mounted using Airvol (Mowiol 4-88; Sigma Aldrich) prior to imaging. Immunofluorescence images were obtained using a Zeiss LSM 800 Confocal Microscope from the Schulich Imaging Core Facility (University of Western Ontario) with a Plan-Apochromat 63x/1.40 Oil DIC objective (Carl Zeiss, Oberkochen, Germany). Representative brightfield images captured with a Nikon Eclipse TE2000-S inverted microscope equipped with RT slider SPOT (HRD 100-NIK) camera are shown.

### Clariom^TM^ S expression profiling and analysis

RNA from 3 biological replicates of A375-P parental cells and PANX1 knockdown A375-P cells were used for Clariom^TM^ S expression profiling at the Genetic and Molecular Epidemiology Laboratory at McMaster University. Expression profiling data were analyzed using TAC version 4.0.2.15 (ThermoFisher Scientific) and RStudio Cloud (https://www.rstudio.com/products/cloud/) version 4.1.2 (RStudio Team, 2022) software to find differentially expressed genes between control and PANX1 knockdown A375-P cells. Differential expression was represented by a gene-level fold change of minimum ±2 and p < 0.05 in PANX1 knockdown samples compared to controls. STRING plots (https://string-db.org/) were generated using the first 100 highest ranked differentially expressed genes showing ±2-fold expression changes, setting the organism to *Homo sapiens*, and kmeans clustering^33,34^.

## Statistical analyses

All data are representative of at least three independent experiments conducted with three technical replicates, unless otherwise stated in the figure legends. Statistical analyses were performed using GraphPad Prism software (version 8.0; San Diego, CA). Error bars indicate mean ± standard error. Details of statistical analysis are provided in the figure legends.

## Results

### Pannexin 1 mRNA correlates with mRNA expression of key components in the Hippo pathway

Gene expression correlations could potentially serve as an indicators of cancer development^35^. To determine whether there is a functional correlation between PANX1 and the Hippo effectors YAP/TAZ in melanoma cells, we initially analyzed a panel of melanoma biopsies from The Cancer Genome Atlas (TCGA, PanCancer Atlas). We observed a weak, but statistically significant, positive correlation between PANX1 mRNA and that of YAP and TAZ in 448 patients with malignant melanoma (Figure 1A and B). IQGAP1 directly binds YAP^36^ and is a scaffold for Hippo components ^22^. Our analysis revealed that PANX1 mRNA exhibits a significant positive correlation with IQGAP1 mRNA in melanoma biopsies (Figure 1C). To investigate whether these correlations are restricted to melanoma or apply to other types of cancers, we analyzed TCGA data of 1108 biopsies taken from patients with breast carcinoma. Similar to melanoma, mRNA expression of PANX1 shows a significant positive correlation with that of YAP, TAZ and IQGAP1 (Figure 1D-F). Overall, these data indicate that in melanoma and breast carcinoma biopsies, PANX1 mRNA expression correlates positively with components of the Hippo pathway, suggesting a potential functional relationship or co-regulation between the PANX1 and core Hippo effectors.

**Fig. 1.**
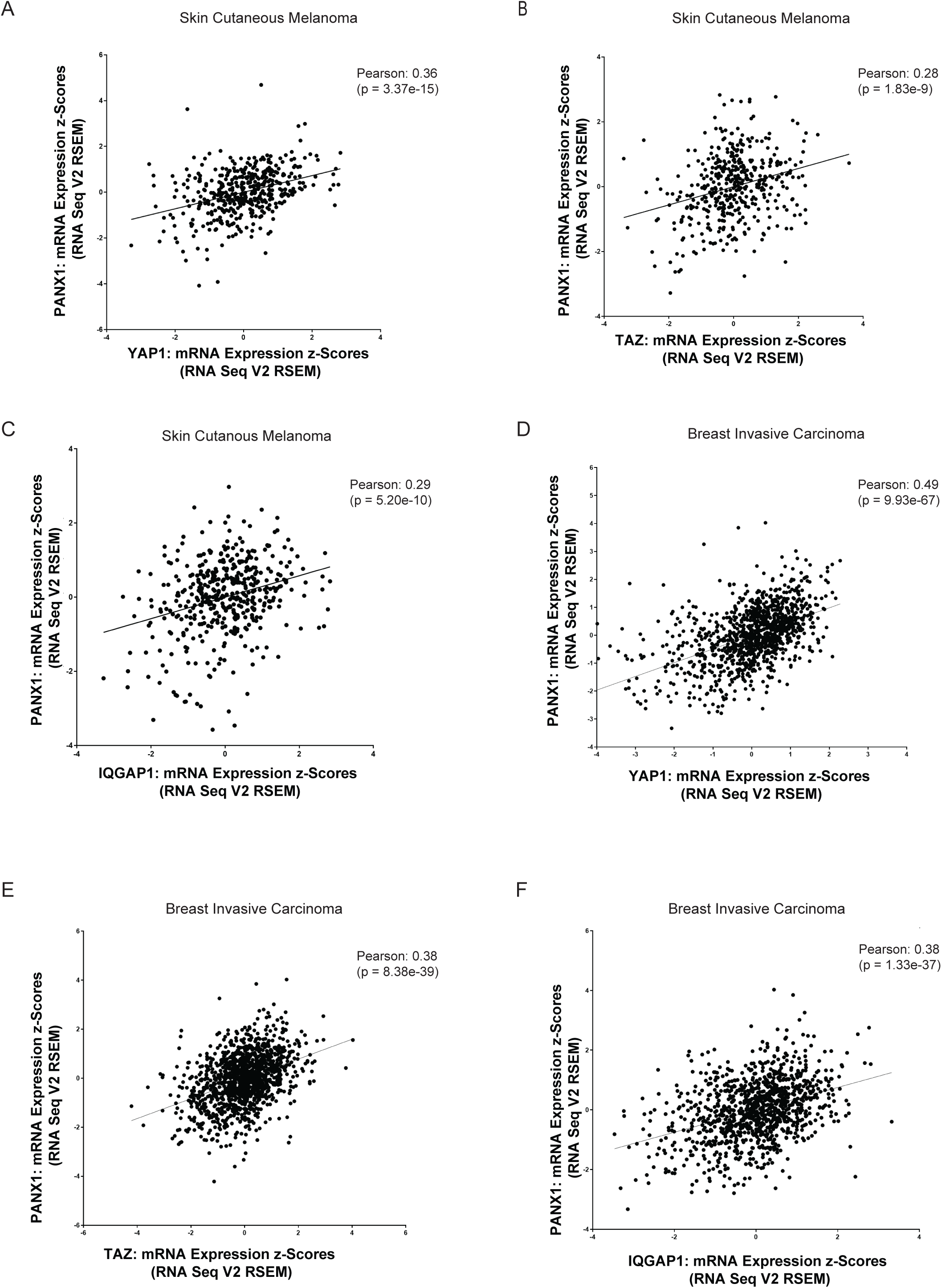
Pannexin 1 mRNA correlates with the mRNA of key components in the Hippo pathway. Analysis of correlation between PANX1 mRNA and YAP, TAZ and IQGAP1 mRNA expression in melanoma (448 samples, A-C) and breast carcinoma (1108 biopsies, D-F) neoplasms in The Cancer Genome Atlas (TCGA) database, revealed that there is a modest yet significant correlation between the PANX1 and IQGAP1, YAP and TAZ mRNA expression levels in melanoma (A-C) and breast carcinoma (D-F).

### Pannexin 1 expression is increased in invasive melanoma cells

Next, we analyzed the abundance of PANX1 in two melanoma cell lines, which have varying invasive capacities. PANX1 protein expression in the more invasive A375-MA2 cells was higher than in the non-invasive A375-P cell line (Figure 2 A and B, *left panel*). The protein level of YAP was also significantly higher in A375-MA2 than A375-P cells (Figure 2B, *right panel*). We evaluated whether the increased YAP expression leads to increased YAP function. CTGF is a YAP-target gene that is widely used as a readout of YAP co-transcriptional function^37^. Quantitative PCR analysis revealed that CTGF mRNA is significantly higher in A375-MA2 cells than in A375-P cells (Figure 2C), indicating that increased YAP abundance in A375-MA2 cells correlates with increased YAP co-transcriptional activity. All together, our data show that PANX1 protein level is increased in the highly invasive melanoma cells and higher amounts of PANX1 correlates with augmented expression and transcriptional function of the Hippo effector, YAP.

**Fig. 2.**
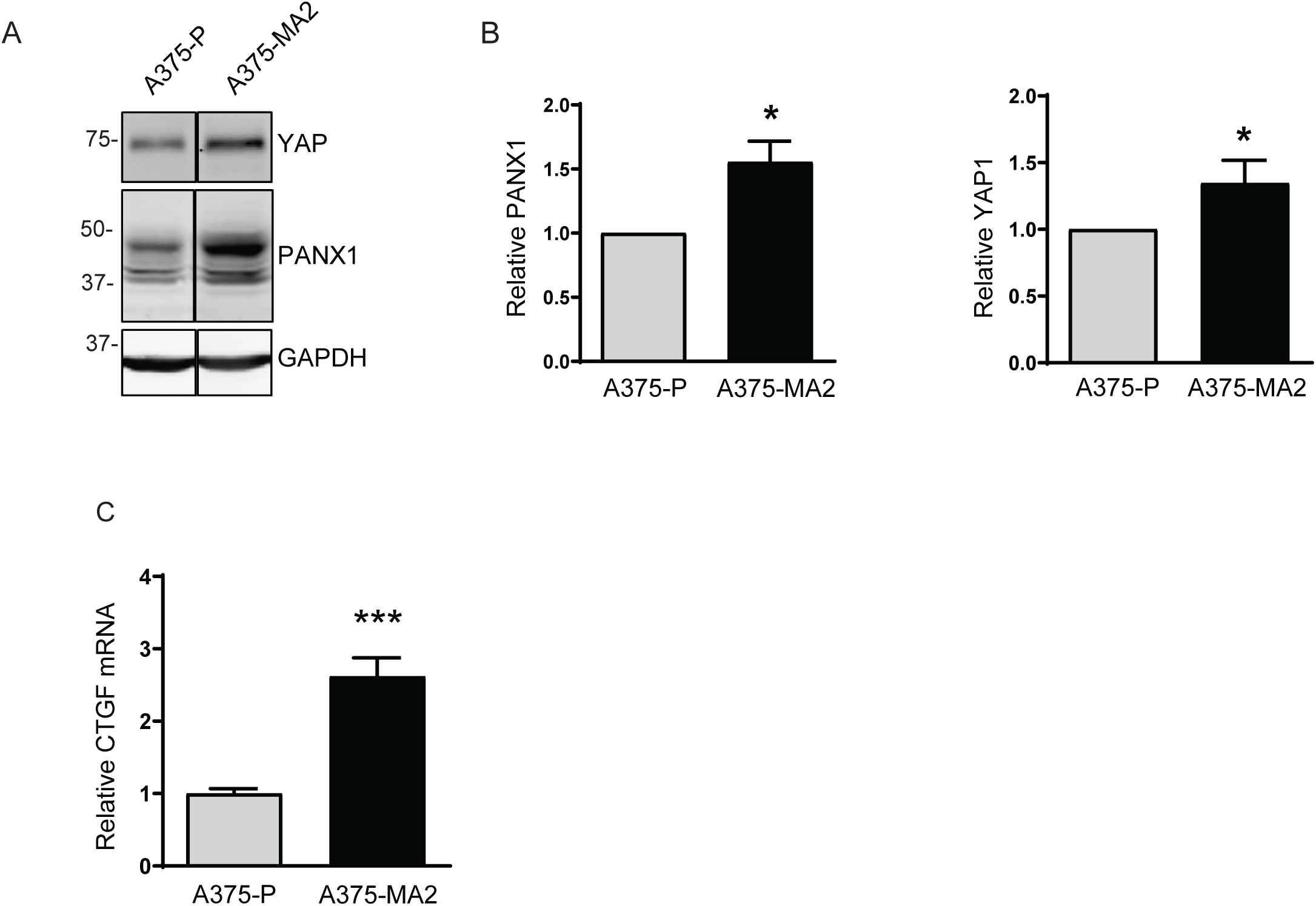
Differential expression of PANX1 and YAP in melanoma cells. (A) Equal amounts of protein lysate from A375-P and A375-MA2 melanoma cells were resolved by SDS-PAGE. Western blotting was performed using antibodies against YAP and PANX1. GAPDH was used as a loading control. Representative Western blots are shown. The lines indicates where irrelevant lanes were removed from the blot. All lanes were scanned at the same intensity. The positions of migration of molecular weight markers are indicated in kDa. (B) The graphs are derived from quantification of Western blots from three or four independent experiments for PANX1 and YAP, respectively, and represent means ± S.E. (error bars) with A375-P cells set as 1. Statistical analysis was conducted using Student’s *t*-test, \**p*<0.05. (C) Quantitative RT-PCR was conducted to measure human CTGF mRNA. The amount of mRNA was corrected to YWHAZ as control in the same sample. mRNA in the A375-P cells were set to 1. The data represent the means ± S.E. (error bars) of three separate experiments, each performed with two technical replicates (*N*=3, *n*=2). Student’s *t* test *** *p*<0.001.

### Endogenous PANX1 associates with endogenous IQGAP1 in melanoma cells

IQGAP1 functions as a scaffold protein in several signaling pathways, including the mitogen-activated protein kinase (MAPK)^38^, phosphoinositide-3-kinase/protein kinase B (PI3K/AKT) cascades^39^ and Hippo signaling ^22^. IQGAP1 binds directly to YAP and suppresses its transcriptional activity^36^. Since PANX1 is predominantly localized in the cytoplasm of melanoma cells (Figure 7A) where IQGAP1 is detected^40^, we investigated whether PANX1 associates with IQGAP1 in melanoma cells. We conducted immunoprecipitation studies in several human melanoma cell lines, including low metastatic A375-P cells as well as the highly metastatic lines A375-MA2 and 131/4-5B1. Immunoprecipitation revealed that endogenous IQGAP1 co-immunoprecipitated with endogenous PANX1 from cell lysates of A375-P (Figure 3A), A375- MA2 (Figure 3B) and 131/4-5B1 (Figure 3C) cells. No IQGAP1 immunoprecipitated with IgG, validating the specificity of the binding. Reciprocally, PANX1 co-immunoprecipitated with endogenous IQGAP1 from 131/4-5B1 cells (Figure 2C, middle panel). The interactions of IQGAP1 with several binding partners, such as calmodulin and B-Raf, are regulated by Calcium^41^. In order to investigate whether Ca^2+^ alters the interaction between IQGAP1 and PANX1, we added either 1 mM Ca^2+^ or 1 mM EGTA to cell lysates before immunoprecipitation. Ca^2+^ did not alter the total amounts of PANX1 or IQGAP1 (Figure 3, Lysate), nor the interaction between the two proteins (Figure 3, IPs). Thus, the interaction between PANX1 and IQGAP1 is not regulated by Ca^2+^. Moreover, we conducted confocal microscopy to study the subcellular domains that PANX1 and IQGAP1 interact. Analysis revealed that PANX1 and IQGAP1 colocalize both at cell border (Figure 3D, black arrow) and intracellular compartments of A375-P cells (Figure 3D, white arrow). Taken together, our data indicate that endogenous PANX1 is in a complex with endogenous IQGAP1 in melanoma cells.

**Fig. 3.**
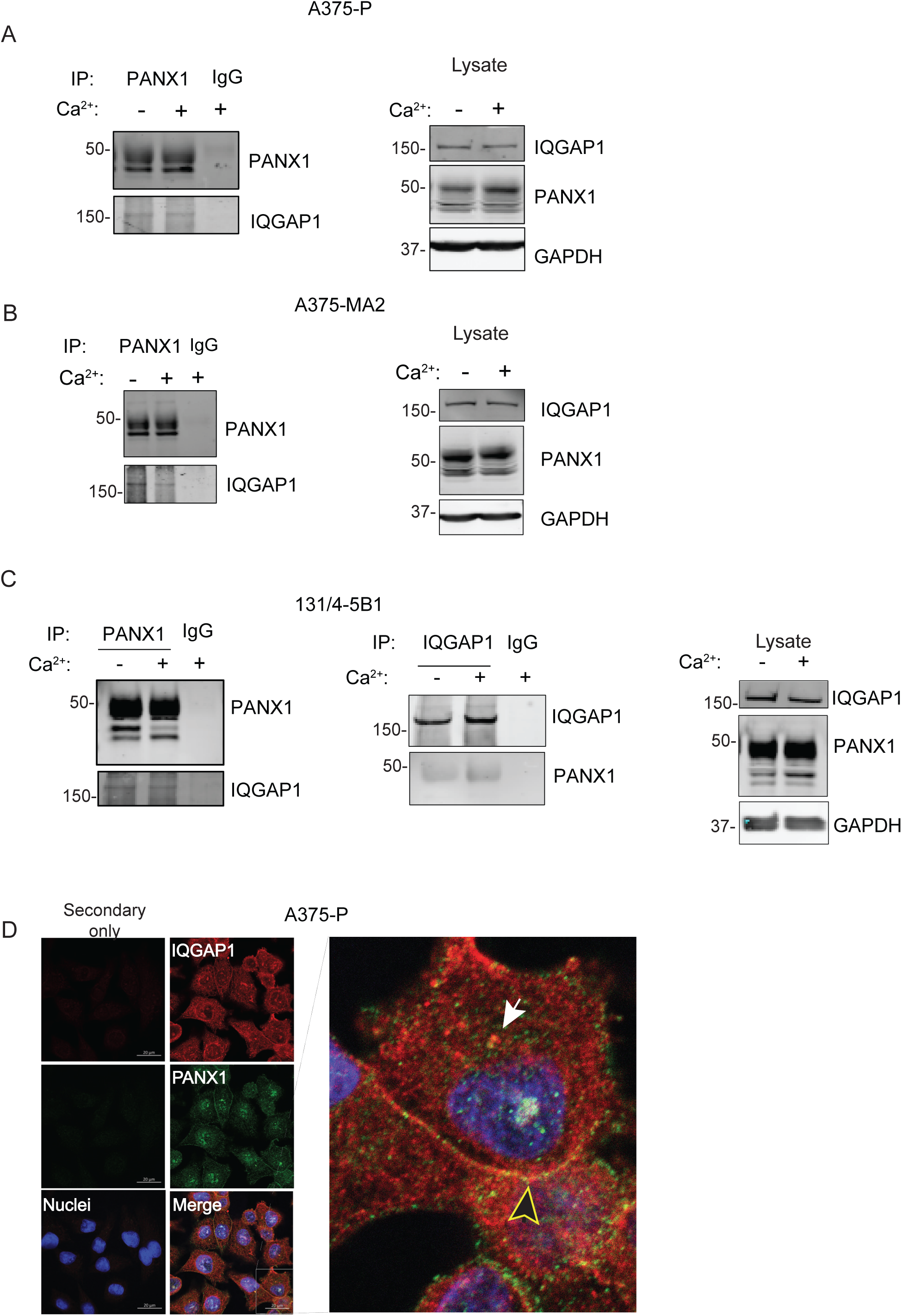
Endogenous PANX1 associates with IQGAP1 in melanoma cells. Equal amounts of protein lysates (1 mg) from A375-P (A), A375-MA2 (B) and 131/4-5B1 (C) cells were obtained with lysis buffer containing 1 mM Ca^2+^ (+) or 1 mM EGTA (-). Lysates were incubated with anti-PANX1 (left panels) or anti-IQGAP1 (C-middle panel) antibodies. Immune complexes were isolated with protein A beads and analysed by SDS-PAGE and Western blotting. Blots were probed with anti-PANX1 and anti-IQGAP1 antibodies. Rabbit or mouse IgG was used as negative controls. Lysates not subjected to IP were also loaded directly onto the gel (*Lysate, right panels*). Anti-GAPDH antibody was used as loading control. All lanes were scanned at the same intensity. The positions of migration of molecular weight markers are indicated in kDa. Data are representative of at least three independent experiments. (D) A375-P cells were fixed and processed for confocal microscopy using antibodies against IQGAP1 (red) and PANX1 (green). DNA was visualized using Hoechst (blue). Inset displays a magnified section of the area outlined by the box. Arrows show areas of colocalization. Scale: 20µm

### IQGAP1 modulates tumorigenic properties in melanoma cells

To initiate analysis of IQGAP1 in the pathogenesis of melanoma, we quantified IQGAP1 protein levels in several human melanoma cell lines from varying stages of disease progression and mutational background (Table 1, Figure 4A).

**Fig. 4.**
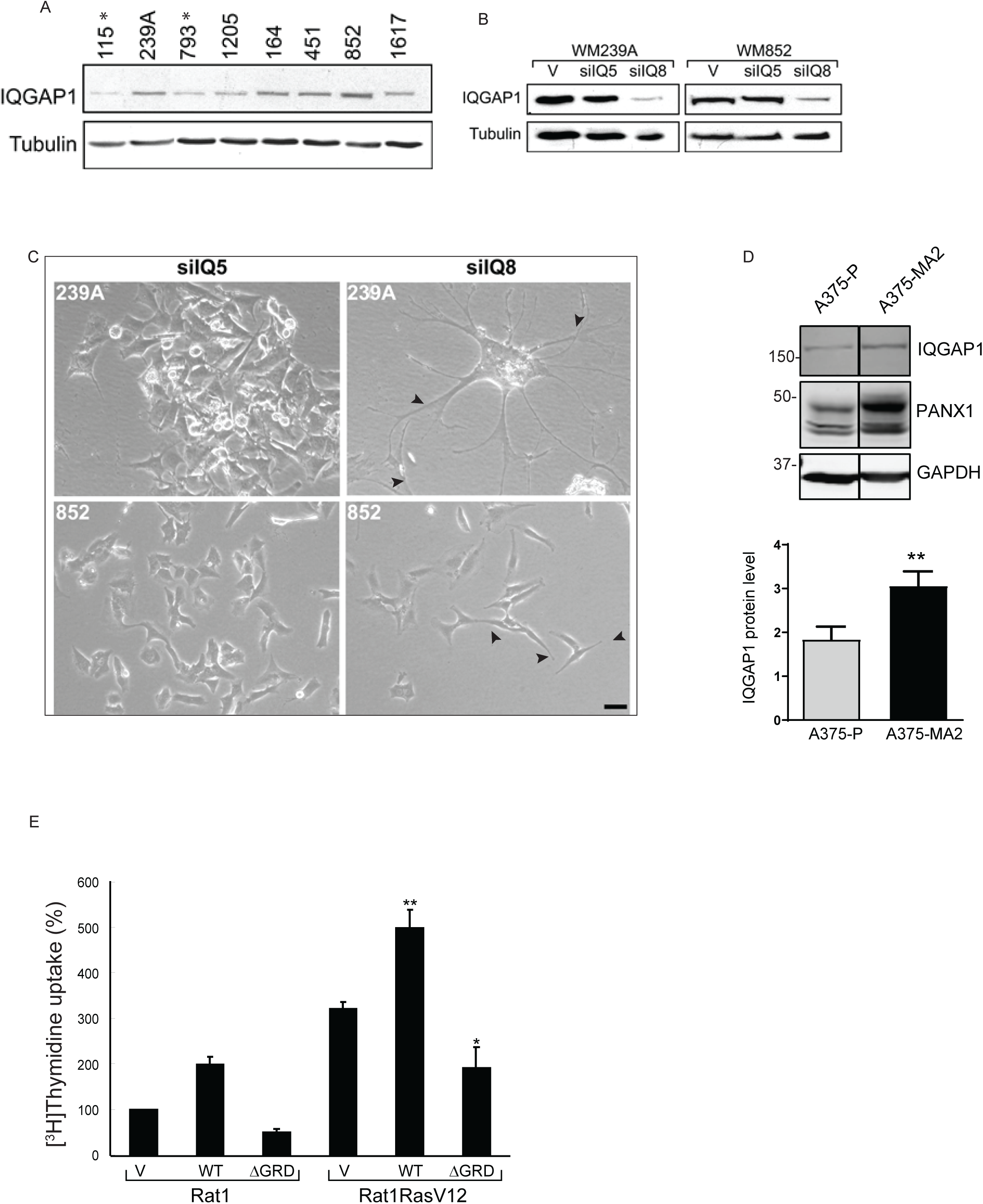
The effect of IQGAP1 on cell proliferation. (A) IQGAP1 in melanoma cell lines. Equal amounts of protein lysates from several human melanoma cell lines (see Table 1) were resolved by Western blotting and probed for IQGAP1. β-tubulin was used as loading control. Data are representative of two independent determinations. Stars are used to denote cell lines that exhibit lower metastatic potential. (B) WM239A and WM852 cells were infected with pSUPER retroviral vector (V) or siRNA constructs siIQ5 or siIQ8. After selecting cells with stable incorporation, equal amounts of cell lysate were processed by Western blotting and probed with anti-IQGAP1 antibodies. β-tubulin was used as loading control. (C) WM239A (upper panels) and WM852 (lower panels) cells, transfected with control (siIQ5, left) or siIQ8 to knockdown IQGAP1 (siIQ8, right), were subjected to microscopy. Arrow heads indicate the areas of dendrites and cell projections. Scale bar, 10 μm. (D) Equal amounts of protein lysates from A375-P and A375-MA2 cells were resolved by SDS-PAGE. Western blotting was performed using antibodies against IQGAP1 and PANX1. GAPDH was used as a loading control. Representative Western blots are shown. The line indicates where irrelevant lanes were removed from the blot. All lanes were scanned at the same intensity. The positions of migration of molecular weight markers are indicated on the left in kDa. The graphs (lower panel) are derived from quantification of Western blots from three independent experiments and represent means ± S.E. (error bars). Statistical analysis was conducted using Student’s *t*-test, \*\**p*<0.01. (E) Rat1 or Rat1RasV12 cells were transiently transfected with either vector (V), wildtype (WT) IQGAP1 or IQGAP1ΔGRD (ΔGRD). After 24 h a [^3^H]thymidine uptake assay was conducted. All assays were performed in quadruplicate. Data are expressed as a percentage of [^3^H]thymidine uptake in vector-transfected Rat1 cells and represent the means ± S.E. (*n*=5 for Rat1, *n*=6 for Rat1V12Ras). * *p*<0.05; \*\**p*<0.001 (significantly different to vector-transfected Rat1RasV12 cells).

**Table 1:**
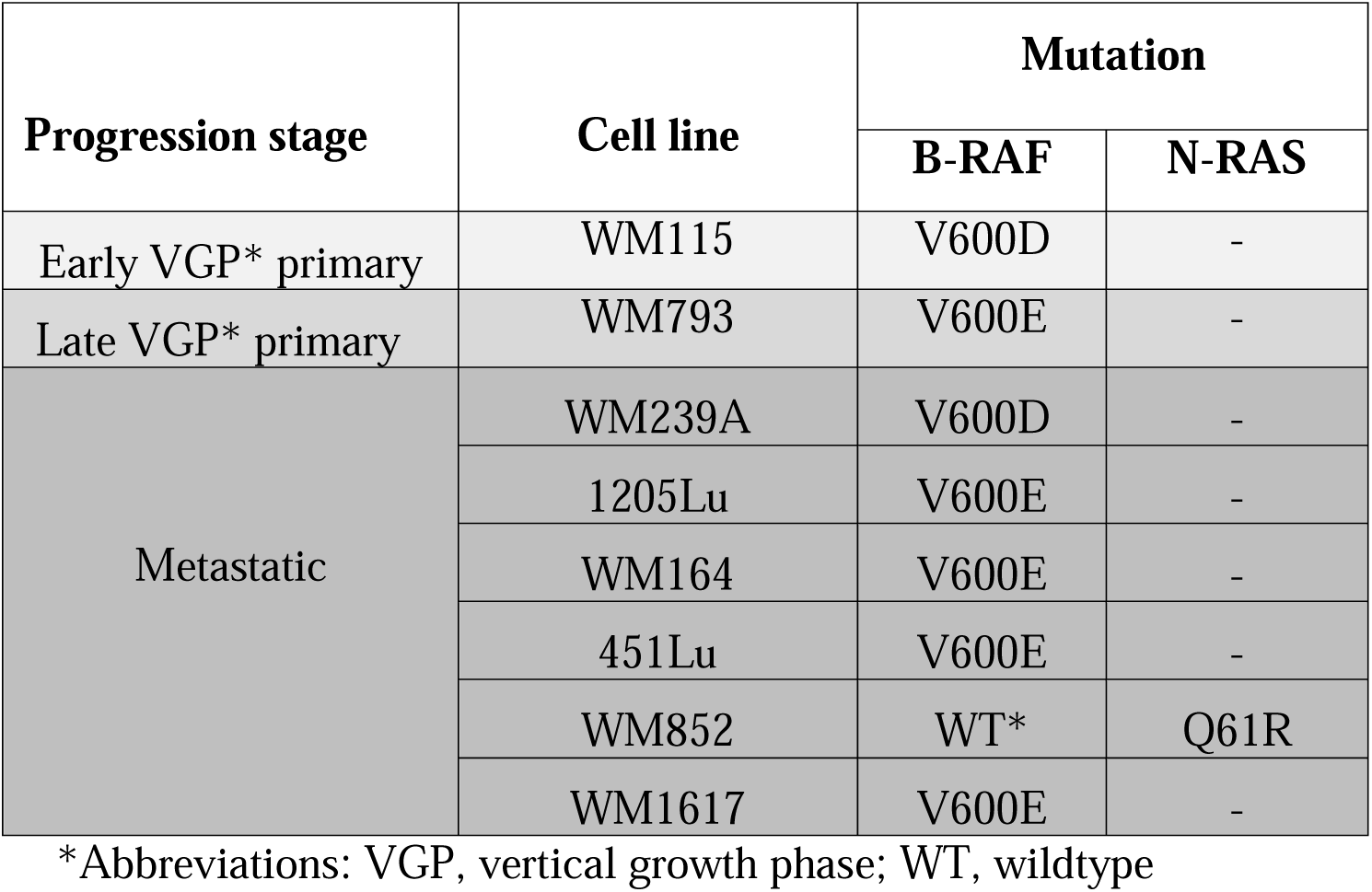
Mutational status and stage of progression of various melanoma cell lines.

| Progression stage | Cell line | Mutation |  |
| --- | --- | --- | --- |
|  |  | B-RAF | N-RAS |
| Early VGP* primary | WM115 | V600D | - |
| Late VGP* primary | WM793 | V600E | - |
| Metastatic | WM239A | V600D | - |
|  | 1205Lu | V600E | - |
|  | WM164 | V600E | - |
|  | 451Lu | V600E | - |
|  | WM852 | WT* | Q61R |
|  | WM1617 | V600E | - |

\*Abbreviations: VGP, vertical growth phase; WT, wildtype

Two features appear from the initial data: i) when we compared early tumorigenic vertical growth phase (VGP) in primary WM115 with metastatic WM239A cells from the same patient, we noticed that IQGAP1 protein concentration tends to be higher in the metastatic lines (Fig. 4A). ii) this observation aligns with findings in WM852 metastatic melanoma cells, which lack BRAF mutations (Fig. 4A). In order to ascertain its role in tumorigenesis, we altered IQGAP1 levels in selected melanoma lines. Using siRNA in a viral vector, we specifically and stably knocked down endogenous IQGAP1 by 75-85% in WM239A and WM852 cells (Fig. 4B). siIQ8 has been well characterized and documented to specifically reduce IQGAP1^31,42^, whereas the oligonucleotide siIQ5, directed against a different region of IQGAP1, does not reduce IQGAP1 expression (Fig. 4B)^31^ and serves as the control. We found that reducing endogenous IQGAP1 levels has a dramatic effect on the morphology of melanoma cells. Compared to cells infected with the non- silencing siIQ5, both WM852 and particularly WM239A cells with IQGAP1 knockdown exhibited a multi-dendritic appearance (Fig. 4C). Over 90% of WM239A cells with IQGAP1 knockdown exhibited the phenotype shown in Fig. 4C. These findings are consistent with differentiation and senescence resulting from the specific reduction in IQGAP1 levels. Overall, our data suggests that IQGAP1 is differentially expressed in melanoma cells and its expression contributes to cell morphology.

A substantial body of evidence suggests that IQGAP1 promotes tumorigenesis^43,44^. We evaluated the expression of IQGAP1 in non-invasive A375-P and metastatic A375-MA2 cell lines. Western blot analysis revealed that the expression of IQGAP1 is significantly higher in invasive A375-MA2 cells than in their non-invasive parental line (Figure 4D).

To investigate the molecular mechanism by which IQGAP1 regulates tumorigenesis, we assessed the effect of IQGAP1 on cell proliferation. Transient overexpression of wildtype IQGAP1 enhanced proliferation of Rat1 fibroblasts by 2-fold (Fig. 4E). By contrast, IQGAP1ΔGRD, which is a dominant negative IQGAP1 construct^45^, reduced DNA synthesis revealed by reduced thymidine uptake (Fig. 4E). Constitutively active RasV12 promotes cell proliferation through several pathways, including Raf/MEK/ERK^46^. Stable overexpression of RasV12 increased Rat1 cell proliferation by 3-fold (Fig. 4E). Transient overexpression of wildtype IQGAP1 further augmented this effect. Importantly, transfection of the dominant negative IQGAP1ΔGRD significantly reduced the ability of RasV12 to increase cell proliferation (Figure. 4E). Thus, IQGAP1 participates in cell proliferation, potentially through its crosstalk with Raf/MEK/ERK signaling. Overall, our data indicate that IQGAP1 is differentially expressed in melanoma cells and its expression modulates both cell morphology and cell proliferation.

### PANX1 regulates YAP protein level

To investigate whether PANX1 regulates the abundance of YAP, we used the CRISPR/Cas9 system to knockout *PANX1* from breast carcinoma cells. Depletion of PANX1 from Hs578T cells significantly decreases the protein level of YAP by 45% (Figure 5A) and the mRNA of YAP target gene, CTGF, by 80% (Figure 5B).

**Fig. 5.**
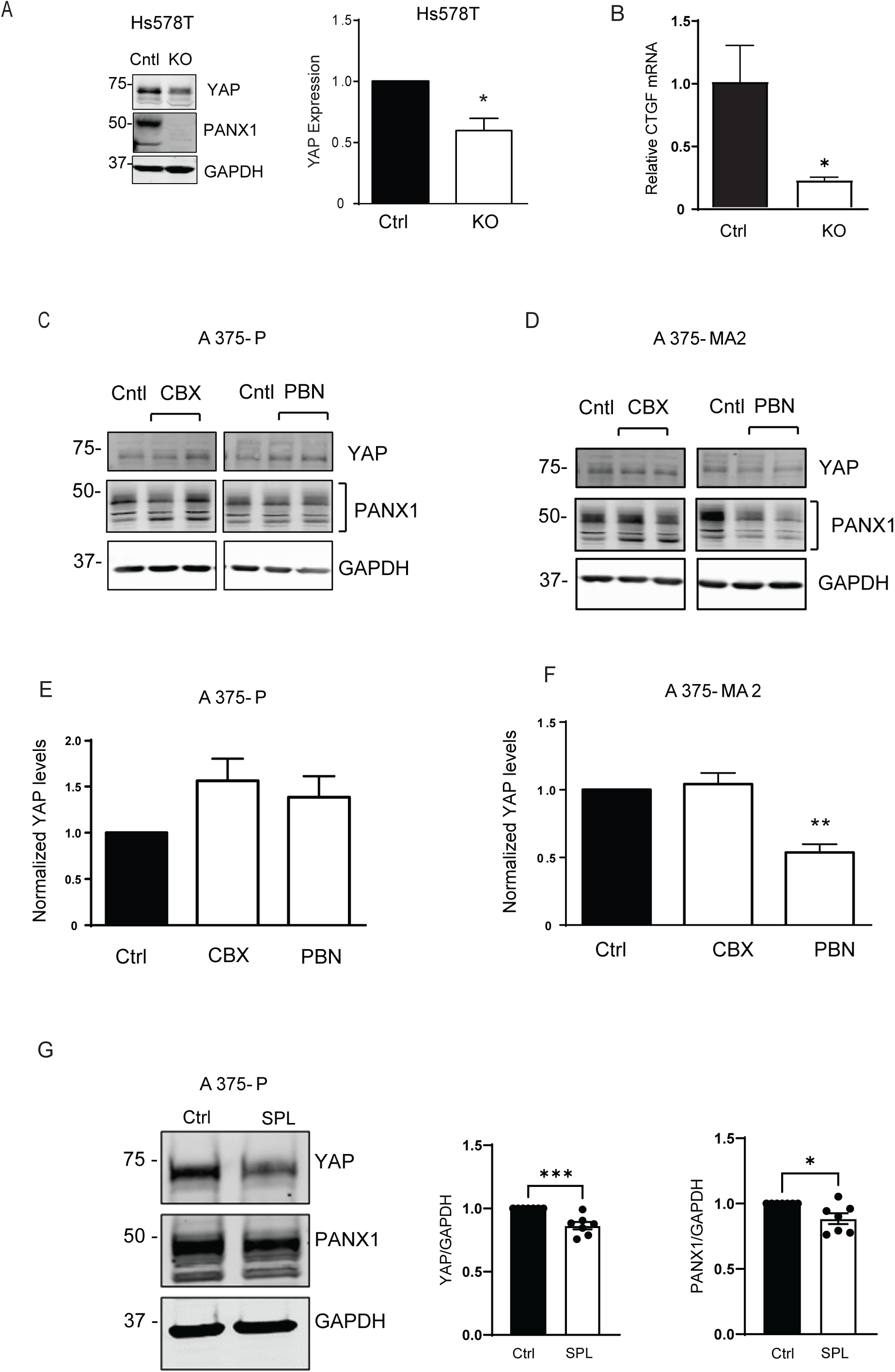
PANX1 regulates YAP protein levels. (A) Equal amounts of protein lysates from Hs578T control (Ctrl) and *PANX1* KO cells were resolved by SDS-PAGE. Western blotting was performed using antibodies against PANX1 and YAP. GAPDH was used as a loading control. The graph (right panel) is derived from quantification of Western blots from three independent experiments. Data represent the mean ± S.E. (error bars) with control cells set to 1. Statistical analysis was conducted using Student’s *t* test \**p*<0.05. (B) Quantitative RT-PCR was conducted to measure human CTGF mRNA from Hs578T control (Ctrl) and PANX1 KO cells. The amount of mRNA was corrected to YWHAZ as control in the same sample. mRNA in the control (Ctrl) cells were set to 1. Data represent the mean ± S.E. (error bars) of three separate experiments, each performed with two technical replicates (*N*=3, *n*=2). Student’s *t* test \**p*<0.05. A375-P (D) or A375-MA2 (E) cells were incubated with 100 μM carbenoxolone (CBX) or 1 mM probenecid (PBN) for 72 h. HBSS was used as vehicle control (Ctrl). Equal amounts of protein lysate were resolved by SDS-PAGE. Western blotting was performed using the antibodies indicated. GAPDH was used as a loading control. The graphs (lower panels) are derived from quantification of Western blots from three independent experiments and represent means ± S.E. (error bars) with control (Ctrl) set as 1. Statistical analysis was conducted using ANOVA \*\**p*<0.01. (G) A375-P cells were incubated with 10 μM spironolactone (SPL) for 72 h. DMSO was used as vehicle control (Ctrl). Equal amounts of protein lysate were resolved by SDS-PAGE. Western blotting was performed using the antibodies indicated. GAPDH was used as a loading control. The graphs are derived from quantification of Western blots from seven independent experiments and represent means ± S.E. (error bars) with control (Ctrl) set as 1. Statistical analysis was conducted using Student’s *t* test \**p*<0.05, \*\*\**p*<0.001.

The expression and function of YAP are highly dependent on the specific cell type and tissue context ^47^. To determine if the interaction between PANX1 and YAP is cell type or cancer context specific, we investigated the levels of total and phospho-YAP in normal liver and skin tissue. We compared the amounts of YAP in livers from WT and PANX1 knockout mice (Figure 7B). Analogous to our findings in breast carcinoma lines, depletion of PANX1 significantly decreases in the abundance of YAP in mouse liver (Figure 7B). Increased YAP phosphorylation induces cytoplasmic retention as well as YAP proteasomal degradation^11^, which reduces YAP transcriptional activity^11^. Depletion of PANX1 from the skin of *PANX1* and *PANX3* double knockout (dKO) mice significantly increases the ratio of phosphorylated YAP^S127^ (pYAP) to total YAP protein (Figure 7C), suggestive of reduced YAP activity. Although tissue-specific mechanisms may play role in the regulation of YAP expression by PANX1, our data indicate that PANX1 modulates YAP levels both in cells and tissues.

Selective FDA-approved PANX1 inhibitors, such as spironolactone, carbenoxolone and probenecid, can effectively interfere with PANX1 function in cells ^48–50^. Although the mechanisms of action of these drugs are not fully elucidated, some of these inhibitors can interfere with binding of PANX1 to intracellular signaling proteins, an effect which may be independent of PANX1 channel function^51^. To investigate the effect of PANX1 inhibition on YAP protein levels, we incubated A375-P and A375-MA2 melanoma cells with either carbenoxolone (CBX)^50^ or probenecid (PBN)^49^ as selective PANX1 chemical inhibitors (Figure 5 C and D). Neither carbenoxolone nor probenecid changed YAP protein levels in non-invasive A375-P melanoma cells (Figure 5E). In contrast, probenecid but not carbenoxolone, significantly reduced the abundance of YAP in the metastatic A375-MA2 cell line (Figure 5F). Spironolactone (SPL) has been recently identified as a selective inhibitor of PANX1^48^. We evaluated whether spironolactone could reduce YAP levels in melanoma cells. Consistent with our findings using probenecid in other cell lines, spironolactone significantly reduced the abundance of YAP in A375-P melanoma cells (Figure 5G). We have previously reported that prolonged exposure to blockers decreases PANX1 levels in melanoma cells^52^. In agreement with this observation, incubating melanoma cells with spironolactone resulted in a mild yet significant reduction in total PANX1 levels (Figure 5G). Therefore, the impact of PANX1 blockers on YAP reduction could be attributed to the decreased abundance of PANX1 and/or inhibition of its channel function. Taken together, chemical inhibition of PANX1 using probenecid and spironolactone, two selective PANX1 inhibitors, effectively decreases the amount of YAP in melanoma cells.

### Clariom™ S analysis of PANX1 knockdown melanoma cells reveals the pathways related to proliferation, adhesion and epithelial mesenchymal transition (EMT)

To evaluate the overall impact of PANX1 loss on cell signaling pathways, Clariom™ S transcriptome analysis was conducted on control and PANX1 KD cells. Principal component analysis (PCA) mapping showed clear separation between the cluster of samples, indicating similar gene expression between similar samples (Figure 6A). Raw data were provided in a ranked list of upregulated to downregulated genes for 21,448 potential candidates when comparing control to PANX1 knockdown cells. Compared to control cells, 424 candidate genes were significantly upregulated or downregulated in PANX1 knockdown samples. We generated a heatmap based on the mRNA alterations of the top 30 lead hits (Figure 6B), revealing a differentially expressed sets of genes between control and PANX1 KD samples. Kyoto Encyclopedia of Genes and Genomes (KEGG) pathway analysis shows the Hippo pathway (code:hsa04390) listed as one of the major pathways affected by PANX1 depletion with a strength of 1.2 and false discovery rate of 0.0043. String pathway analysis revealed that genes related to proliferation and cell adhesion constitute the main altered cluster between control and PANX1 KD samples (Figure 6C). Consistent with our observations (Figure 5B), knocking down PANX1 significantly reduced the YAP target, CTGF mRNA, in A375-P cells (Figure 6D). Additionally, knocking down PANX1 significantly decreases the mRNA of neuroglin1 (NRG1), β-galactoside binding protein, galectin-3 (LGALS3) and metallothionein (MT1E) (Figure 6D), all of which are involved in melanoma pathogenesis^53,54,55^. Moreover, the expression of BARX1, a member of the Bar-like homeobox gene family that potentiates cell proliferation, differentiation, immune responses, and tumorigenesis, is increased in PANX1-deficient cells (Figure 6D). Overall, our Clariom™ S analysis offers a comprehensive insight into the proteins that undergo alterations in the absence of PANX1, shedding light on the potential influence of PANX1 on melanoma progression.

**Fig. 6.**
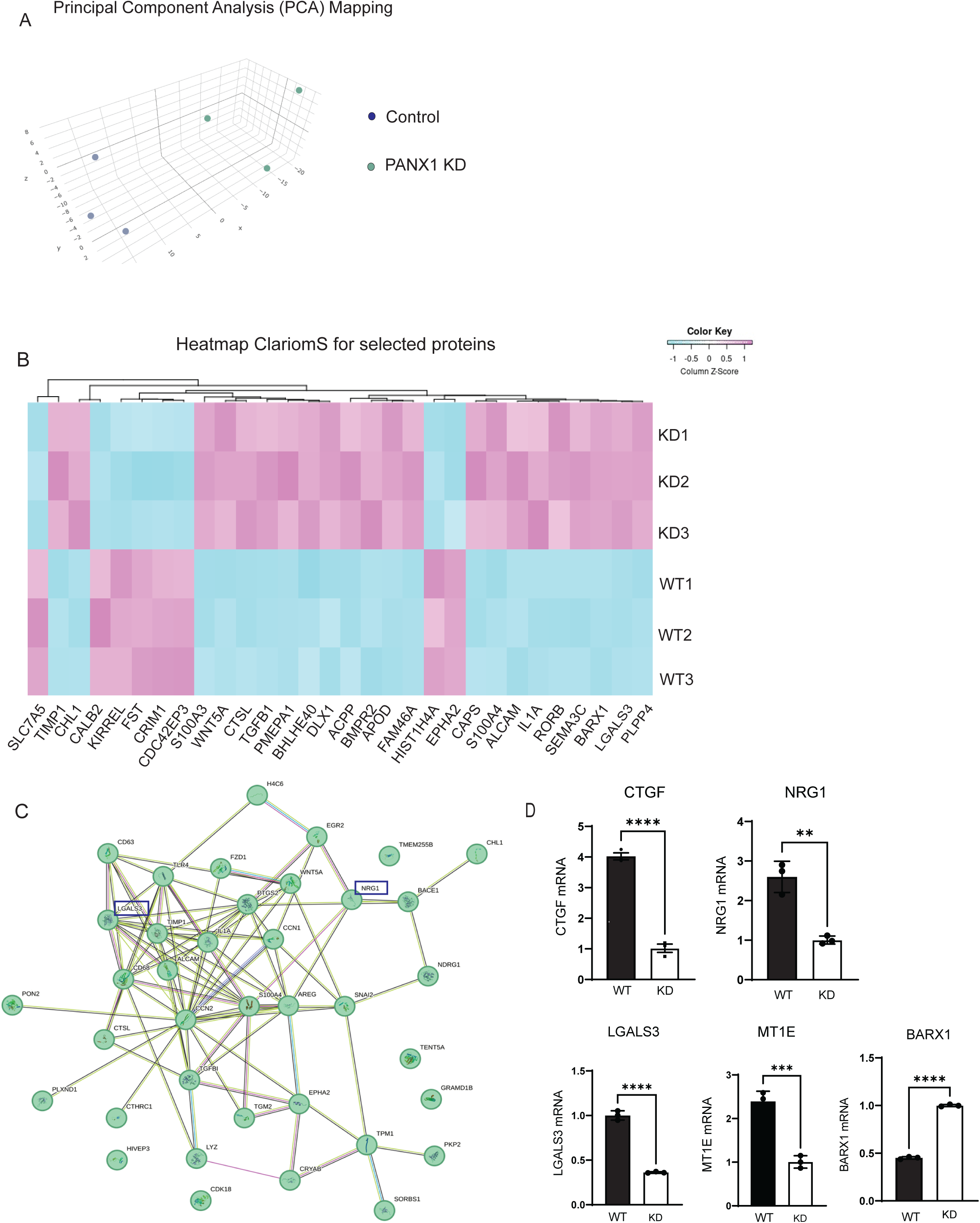
Transcriptome analysis of control shRNA and PANX1 knockdown cells isolated 424 candidate genes. mRNA was isolated from PANX1 knockdown (*N*=3) and control cells (*N*=3) and used for transcriptome analysis. (A) Principal component analysis (PCA) of PANX1 control and knockdown (KD) samples show a clear separation between each cell type cluster. (B) Heat map showing the 30 highest ranked differentially expressed genes between WT (WT 1–3) and PANX1 KD (KD 1–3) samples. Individual genes found to have a gene-level fold change of minimum ±2 column Z-score and *p*<0.05 between WT and KO samples. (C) A STRING plot was generated using the first 100 highest ranked differentially expressed genes showing ±2-fold expression changes, organism *Homo sapiens* and kmeans clustering into three clusters. The major cluster is associated with cell proliferation, and KEGG pathway analysis shows Hippo pathway listed as one of the major pathways affected by PANX1 depletion. Boxes show NRG1 and LGALS3 that are discussed in D. (D) Genes that are differentially expressed between A375-P parental cells (WT) and PANX1 knockdown A375-P cells (KD) selected from Clariom™ S and String analysis. The data represent the means ± S.E. (error bars) of at least three separate samples (*N*=3). Statistical analysis was conducted using Student’s t test \*\**p*<0.01, \*\*\**p*<0.001, \*\*\*\**p*<0.001 compared to control samples.

**Figure 7.**
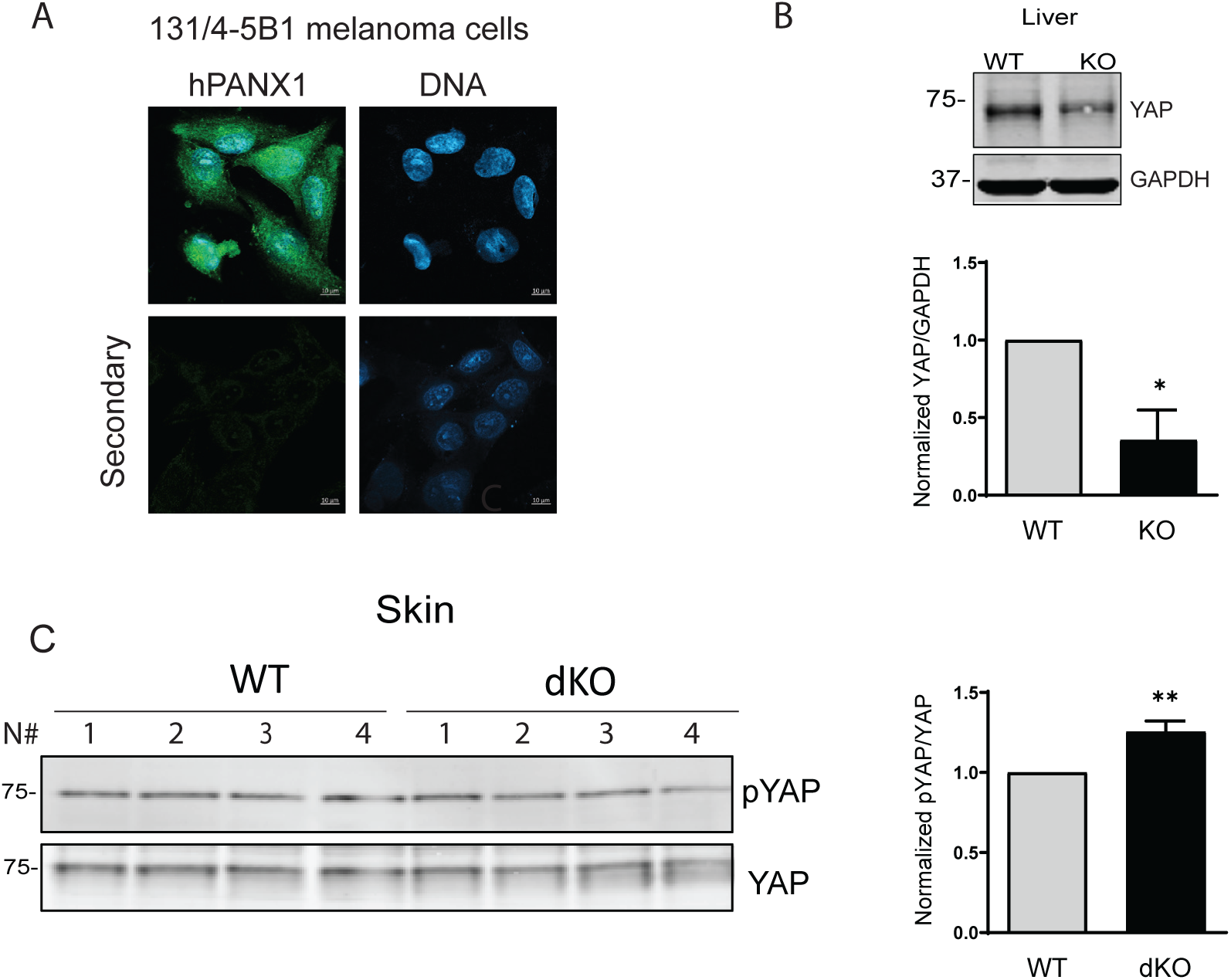
(A) Confocal images of 131/4-5B1 melanoma cells fixed and stained with anti-PANX1 (green). DNA was stained with Hoescht (blue). Samples incubated only with Alexa fluor 488 secondary antibodies were used as control (Secondary). Scale bar, 10μm. (B) Equal amounts of protein lysates from livers of WT and *PANX1* KO mice were resolved by SDS-PAGE. Western blotting was performed using the antibodies indicated. The data represent the mean ± S.E. (error bars) of at least three separate experiments (*N*=3). Statistical analysis was conducted using Student’s t test \**p*<0.05 compared to WT cells set to 1. (C) Equal amounts of protein lysates from the skin of WT and *PANX1/PANX3* double KO (dKO) mice, 4 separate mice (N#1-4) per genotype, were resolved by SDS-PAGE. Western blotting was performed using the antibodies indicated. The data represent the mean ± S.E. (error bars) of 4 separate mice per genotype. Statistical analysis was conducted using Student’s *t test* \*\**p*<0.01 compared to WT cells set to 1.

Gene Set Enrichment Analysis (GSEA) identifies gene subsets with similar biological function, regulation, and chromosomal location^56^. We conducted GSEA analysis on the gene candidates with differential expression between control and PANX1 KD cells (Figure 8A). Five significant gene subsets were identified by GSEA from the Hallmark Gene Database: Hallmark- TNFα Signaling via NF-κB, Hallmark-Hypoxia, Hallmark Epithelial-Mesenchymal Transition, Hallmark-Estrogen Response Late, and Hallmark-IL2 STAT5 signaling. The enrichment score represents the degree to which a gene subset is overrepresented in either extreme (upregulation or downregulation) of the ranked list of genes (Figure 8A). Each gene subset has a leading subset, which is the group of genes that contributes the most to a gene subset’s enrichment score.

**Figure 8.**
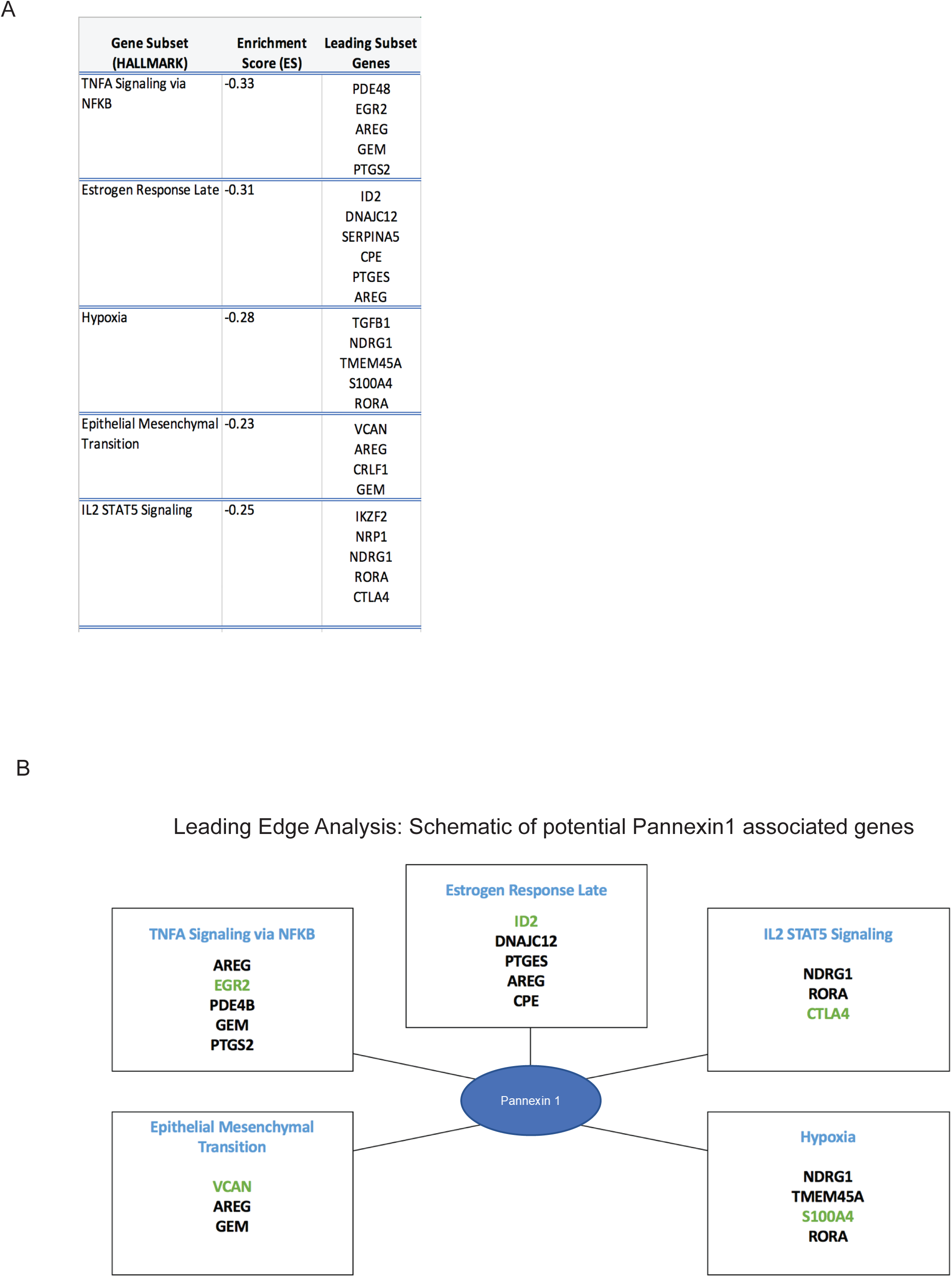
(A) Gene subset enrichment analysis (GSEA) of significant candidate genes from Clariom™ S transcriptome analysis. Five gene subsets (Hallmark-TNFα Signaling via NF-κB, Hallmark-Hypoxia, Hallmark-Epithelial Mesenchymal Transition, Hallmark-Estrogen Response Late, and Hallmark-IL2 STAT5 Signaling) were isolated from 50 common gene subsets registered in the Hallmark Gene Database. Enrichment score (ES) represents the degree to which a gene subset is over-represented on either extreme of the ranked gene list. Leading subset genes contribute the most to the enrichment score. (B) Leading Edge Analysis isolated candidate genes that are most represented in all significant leading gene subsets. Schematic identifies 5 genes (green) for future mRNA and protein analysis due to their potential roles in PANX1-mediated regulation of melanoma: versican (VCAN), early growth response protein 2 (EGR2), DNA-binding protein inhibitor (ID2), cytotoxic T-lymphocyte-associated protein 4 (CTLA4), and S100 calcium binding protein A4 (S100A4) from Hallmark-Epithelial Mesenchymal Transition, Hallmark-TNFα Signaling via NF-κB, Hallmark-Estrogen Response Late, Hallmark-IL2 STAT5 Signaling, and Hallmark-Hypoxia gene subsets, respectively. ± 2**-**fold change, *p*<0.0

We conducted Leading Edge Analysis to pick out relevant genes from each set of significant leading genes. Early growth response protein 2 (EGR2), versican (VCAN), DNA- binding protein inhibitor (ID2), cytotoxic T-lymphocyte-associated protein 4 (CTLA4), and S100 calcium-binding protein A4 (S100A4) were isolated from Hallmark-TNFα Signaling via NF-κB, Hallmark-Epithelial Mesenchymal Transition, Hallmark-Estrogen Response Late, Hallmark-IL2 STAT5 Signaling, and Hallmark-Hypoxia gene subsets, respectively (Figure 8B). Each of these five genes were significantly lower (green) in PANX1 knockdown cells than in control cells and were selected as representative examples of genes known to be involved in regulating melanoma progression^57–61^.

## Discussion

PANX1 was initially discovered as a gap junction channel protein^62^. In recent years, an increasing body of evidence suggests that PANX1 also functions as a signaling protein, modulating pathways such as Wnt signaling^52^, which is a role that is not necessarily tied to its function as a channel. Moreover, PANX1 exhibits intracellular localization in several cancer cell lines, including, but not limited to, colon carcinoma^63^, melanoma^52^ and non-small lung carcinoma cells^64^. The predominant intracellular localization of PANX1 does not conform to the classic channel distribution at the cell border, further indicating potential intracellular roles for PANX1. Here we demonstrate in melanoma cells that endogenous PANX1 is in a complex with IQGAP1, a scaffold protein in the Hippo signaling pathway^22^. Further, we demonstrate that cells with higher expression of PANX1 exhibit increased expression of IQGAP1 and YAP. We found that pharmacological inhibition or depletion of PANX1 reduces the abundance of YAP in melanoma and breast carcinoma cells. Our findings provide insight into the potential role of PANX1 in YAP-mediated cellular processes, such as proliferation, migration, and invasion during melanoma progression.

The Hippo pathway and Wnt signaling are major inducers of transcriptional reprogramming of melanoma cells from a proliferative to invasive mesenchymal phenotype^65^. We previously reported that knocking down PANX1 reduces the tumorigenic properties of melanoma cells and reverts them to a more melanocytic phenotype^20,21^. However, the mechanism(s) through which PANX1 regulates melanoma progression has not been fully elucidated. We recently reported a previously unrecognized crosstalk between PANX1 and Wnt signaling^22^. Importantly PANX1 directly binds to β-catenin, a key transcriptional factor of Wnt signaling^52^. In this study, we show that endogenous PANX1 forms a complex with the scaffold protein IQGAP1. IQGAP1 binds directly to both β-catenin^66–69^ and YAP^36^ and modulates their transcriptional activity. Thus, it is likely that PANX1 participates in the crosstalk between the Wnt and Hippo pathways during melanoma progression.

An important cellular process that both PANX1^70^ and IQGAP1^71,72^ participate in is actin- remodeling. IQGAP1 directly binds to actin^26^ and interacts with several actin-related proteins, including but not limited to adenomatous polyposis coli (APC) and small GTPases, Cdc42 and Rac1^73,74^, and silencing IQGAP1 significantly reduces cell migration^31^. Notably, PANX1 also interacts directly with actin and actin-binding proteins, such as ARP2/3^75^, and silencing PANX1 significantly attenuates migration of melanoma cells ^21^. The actin cytoskeleton is a major regulator of YAP subcellular localization and function^76^. Importantly, actin remodeling induces resistance to the BRAF inhibitor, vemurafenib, by modulating YAP^77^. The impact of the interaction between PANX1 and IQGAP1 and the potential impact on both actin remodeling and YAP subcellular localization should be investigated in future studies.

Overexpression of YAP promotes tumor growth/metastasis in several cancer cells^78,79^ including melanoma^80^, and leads to a distinct transcriptional profile that spans several hallmarks of cancer^81^. Increased activity of the YAP/TAZ pathway has been linked to worse survival in patients^82,83^ and inhibition of YAP/TAZ has been suggested as a potential therapeutic strategy^84^. Our data indicates that PANX1 regulates the abundance of YAP in melanoma and breast carcinoma cells. Our data from the skin tissue of the dKO mice, where PANX3 cannot compensate for the loss of PANX1, along with our findings from the liver of PANX1 KO mice demonstrate that in a more complex context, PANX1 is important for the regulation of YAP levels.

In our pathway analysis, the gene cluster related to proliferation was altered in the absence of PANX1, which could potentially be due to the reduced YAP-mediated proliferation in cells^12^. YAP also enhances migration of melanoma cells via transcriptional regulation of the ARPC5 subunit of the actin nucleating ARP2/3 complex^85^. Notably, ARPC5 is regulated by Ca^2+^/calmodulin, and in melanoma cells, PANX1 forms a complex with calmodulin^52^. The role of Ca^2+^/calmodulin in regulating Hippo signaling is an emerging field of investigation^86^. Given the role of PANX1 in mediating Ca^2+^ release and the interaction between PANX1 and calmodulin in melanoma cells, it is plausible that PANX1 may regulate YAP via Ca^2+^-mediated signaling.

Our Clariom™ S analysis revealed that several oncogenes, including CTGF, NRG1, LGALS3 and MT1E, exhibit reduced transcription in the absence of PANX1. NRG1 is a direct ligand for ErbB tyrosine kinase receptors and altered NRG1 signaling has been reported in malignant melanoma^53^. LGALS3 is increased in patients with metastatic melanoma and silencing galectin-3 in metastatic melanoma cells reduces tumor growth and metastasis^54^. Moreover, MT1E overexpression is a prognostic factor in invasive melanoma^55^. These findings suggest that suppressing PANX1-mediated signaling may protect against melanoma progression.

High expression of YAP/TAZ in malignant melanoma correlates positively with tumor thickness, invasion depth grade, and lymph node metastasis^83^. YAP is reported to induce anoikis resistance in melanoma^13^, and knocking down YAP significantly decreases the number and size of lung metastasis in a mouse xenograft model^13^. Additionally, 50% of melanomas exhibit BRAF mutations and YAP causes BRAF inhibitor resistance in melanoma by inducing T cell expression of programmed death ligand 1 and T cell exhaustion^87^ . Therefore, inhibition of YAP/TAZ offers a potential therapeutic approach in treatment strategies against malignant melanoma.

Pharmacological inhibition of YAP/TAZ by the benzoporphyrin verteporfin, which inhibits YAP/TEAD transcriptional activity, has gained attention in clinical practice because of its effect on inhibiting cell growth in several cancers such as glioma^88^ and glioblastoma^89^, where it is currently in phase I/II clinical trial^90^. While *in vitro* studies show that verteporfin inhibits the growth of certain melanoma cell types, such as BAP1-positive uveal melanoma cells, it is not effective in suppressing the growth of BAP1-negative cells and conjunctival melanoma^91^. Therefore, it is necessary to explore alternative methods of suppressing YAP/TAZ in melanoma. We have previously shown that long term exposure to PANX1 inhibitors will influence PANX1 stability/abundance in cells about 20-40%^52^. Our data indicates that repurposed FDA-approved PANX1 inhibitor, probenecid and spironolactone, decrease the abundance of YAP in malignant melanoma cell lines. Collectively, PANX1 inhibitors, probenecid and spironolactone, can potentially be effective in targeting YAP-mediated signaling as an adjuvant therapeutic intervention in melanoma.

## Author contributions

S.S.: Study design and coordination, performed experiments, data analysis, supervision, original manuscript draft and editing.

K.H.: performed experiments, original manuscript draft editing. C.Z.: performed experiments.

M.K.: performed computational analysis. M.B.: performed computational analysis,

BO’D: performed experiments, original manuscript draft editing

B.W: performed experiments, original manuscript draft editing. Z.L: performed experiments.

D.J: performed experiments, manuscript editing S.E.L: performed experiments, manuscript editing. M.H: performed experiments, data analysis.

L.D: funding acquisition, manuscript editing

D.B.S: conceptual study design, supervision, funding acquisition, manuscript editing S.P.: conceptual study design, supervision, funding acquisition, manuscript editing

## Acknowledgments

This study was supported by a Canadian Institutes of Health Research CIHR Project grant to S. P. (FRN 153112). The authors thank the many wonderful colleagues in the Penuela and Sacks laboratories who have contributed to studies investigating pannexins and the Hippo pathway. We thank Lorraine Jadeski and Jian Guo for initial experiments on melanoma cells. Z. L. and D. B. S. were funded by the Intramural Research Program of the National Institutes of Health.

## Data Availability Statement

The data that support the findings of this study are available in the Materials and Methods, Results, of this article. All the raw data are available upon request.

