## Supplementary figures and images for "Pannexin 1 crosstalk with the Hippo pathway in malignant melanoma"

### Suppl Fig 1

Supplementary Figure 1

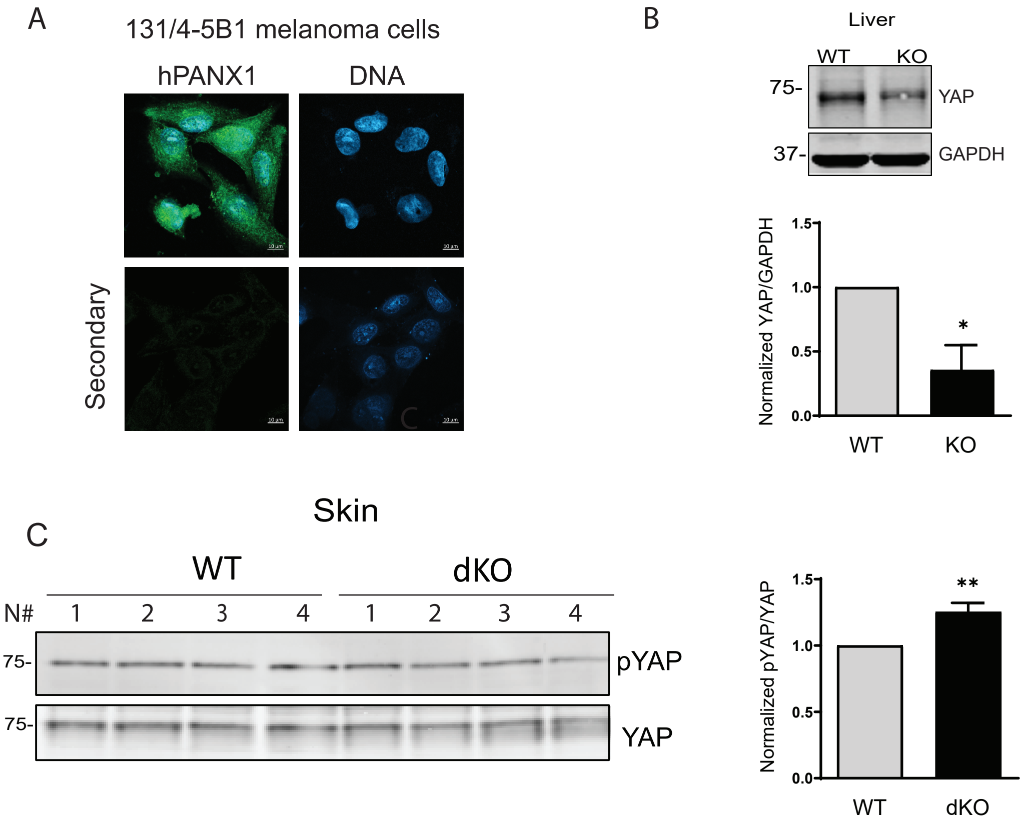
