## Supplementary material for "Pannexin 1 crosstalk with the Hippo pathway in malignant melanoma": Suppl Fig 2

Supplementary Figure 2

A

| Gene Subset (HALLMARK) | Enrichment Score (ES) | Leading Subset Genes |
|---|---|---|
| TNFA Signaling via NFKB | -0.33 | PDE48<br>EGR2<br>AREG<br>GEM<br>PTGS2 |
| Estrogen Response Late | -0.31 | ID2<br>DNAJC12<br>SERPINA5<br>CPE<br>PTGES<br>AREG |
| Hypoxia | -0.28 | TGFB1<br>NDRG1<br>TMEM45A<br>S100A4<br>RORA |
| Epithelial Mesenchymal Transition | -0.23 | VCAN<br>AREG<br>CRLF1<br>GEM |
| IL2 STAT5 Signaling | -0.25 | IKZF2<br>NRP1<br>NDRG1<br>RORA<br>CTLA4 |

B

Leading Edge Analysis: Schemtaic of potential Pannexin1 associated genes

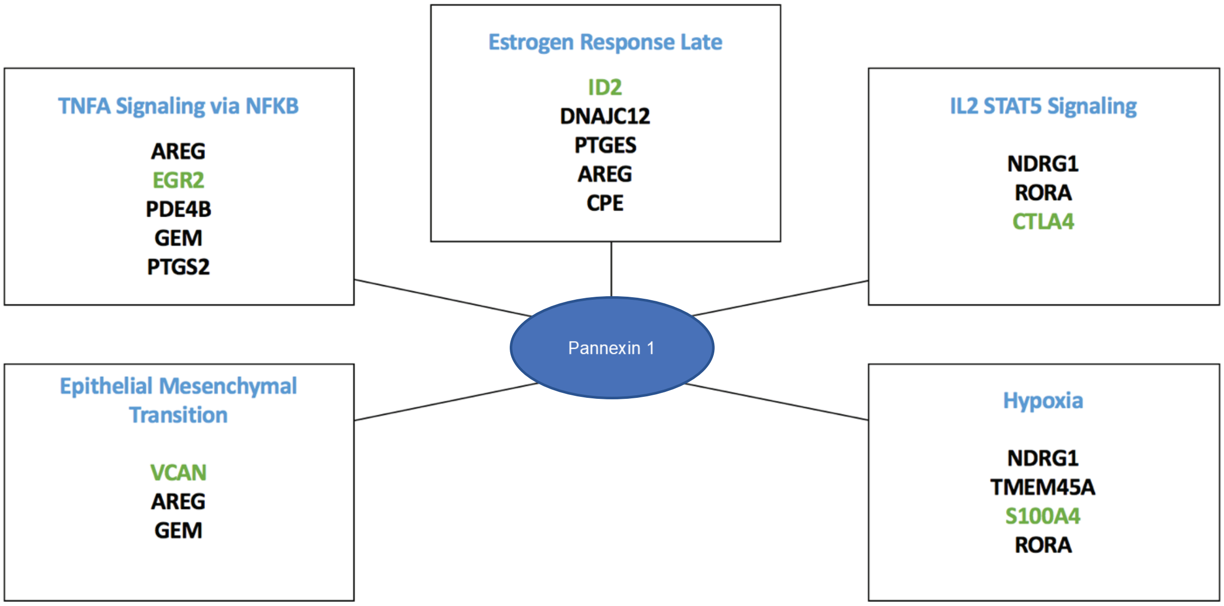
